# A new paradigm for personal protection against ticks: efficacy of spatial repellents to reduce host seeking activities in three major tick species of medical importance

**DOI:** 10.1101/2022.05.17.492268

**Authors:** Eric L. Siegel, Marcos Olivera, Esteban Martinez Roig, Melynda Perry, Andrew Y. Li, Sebastián D’hers, Noel M. Elman, Stephen M. Rich

**Affiliations:** Laboratory of Medical Zoology, Department of Microbiology, University of Massachusetts, Amherst, Massachusetts, USA; Computational Mechanics Center. Instituto Tecnológico de Buenos Aires (ITBA), Ciudad Autónoma de Buenos Aires, Argentina; Textile Materials Evaluation Team, Combat Capabilities Development Command Soldier Center, US Army Garrison-Natick, Natick, Massachusetts, USA; USDA-ARS, Invasive Insect Biocontrol & Behavior Laboratory, Beltsville, Maryland, USA; GearJump Technologies, LLC., Brookline, Massachusetts, USA

**Keywords:** active ingredients (AIs), controlled-release device (CRD), olfaction, pyrethroid, spatial repellency, spatial repellents (SRs), ticks

## Abstract

Addressing the prevalence of tick-borne disease requires robust chemical options as an integral component of Integrative Vector Management (IVM) program. Spatial repellency is a novel concept in tick bite prevention. To date, there is no standard for the evaluation of spatial repellency against ticks, despite the speculated value of volatilized chemicals in control systems. This study reports a novel vertical climb assay that was specifically created for the quantitative evaluation of spatial repellency in ticks. Controlled release devices (CRDs) were used to control the dispersion of multiple Active Ingredients (AIs) transfluthrin, metofluthrin, nootkatone, and DEET against adult females of three medically important tick species: *Dermacentor variabilis, Amblyomma americanum*, and *Ixodes scapulari*s. Results of our study indicate significant associations between AI exposure and changing in tick climbing behavior when compared controls in the absence of the AI, from several perspectives, including changes in tick movement velocity, displacement, detachment, and rate of successful vertical climbing. Metofluthrin and transfluthrin caused strong reductions in host seeking activities against *D. variabilis* and *A. americanum*, while both demonstrated slightly weaker effects against *I. scapularis*. Further work is planned to evaluate spatial repellency in ticks in more natural environments and assess their potential in future tick control programs.

## Introduction

Ticks are the principal arthropod vectors of a variety of human, livestock, and companion animal disease in North America, such as Lyme disease, Anaplasmosis, and Babesiosis [1]. The prevalence of these zoonotic diseases has increased recently due to shifts in host population dynamics, particularly with the white-tailed deer, that affect tick population size [2]. Targeting of live arthropod populations is an important component of Integrative Vector Management (IVM) programs aimed at addressing the climbing incidence of these diseases [3,4]. Interventions used for mitigating risk include source reduction and personal protection. Chemical means of source reduction entail large-scale ground spraying or treatment of reservoir hosts to kill ticks in the environment, while personal protective methods seek to reduce risk to individual humans through smaller-scale, personal chemical application. Repellents can be classified as contact repellents, requiring physical contact with the treated source, or Spatial Repellents (SRs), acting by volatilized AI. With spatial compounds, vapor phase concentration and arthropod inherent sensitivity determine whether repellency will occur over a given space [5]. A new generation of Active Ingredients (AIs) derived from synthetic pyrethroids are often described as SRs and allow for non-contact protection from vectors that cover large distances, such as mosquitoes and biting flies [6]. SRs can maintain their efficacy by sustaining a spatial concentration over time with dispersion control methods. Controlled release devices (CRDs) that modulate SR dispersion therefore play a key role in efficacy [7].

The actions of repellent AIs are based on multiple target biomechanisms, including attraction-inhibition, irritation, and intoxication. The host-seeking behavior and ecology of ticks, however, challenges the applicability of these repellent biomechanisms that are traditionally used to combat more agile, flying arthropods. DEET (*N,N-diethyl-3-methylbenzamide)* is the most prominent commercial arthropod contact repellent and the gold standard against which novel AIs are compared [8]. It deters mosquitoes from landing on a treated surface by interfering with receptors on antennae that would regularly detect host cues, such as heat and carbon dioxide [9]. In ticks, however, DEET has been described to work as a contact irritant – with varied efficacy across multiple species [10]. Other known pyrethroids are also used as contact repellents in arthropod protection. Permethrin is an AI within this class frequently used in ultra-low volume (ULV)-spraying and other source reduction techniques. It repels and kills mosquitoes and other flying arthropods with direct droplet contact [11]. It is also used in personal protection as a clothing treatment for long-lasting tick protection through acaricidal action, with little signs of repellency [12].

The lack of a standardized method for evaluating non-lethal, behavior modifications in ticks stems from an incomplete understanding of tick olfaction at the molecular level and lack of defined actions in the host-seeking and feeding process [13]. Ticks are relatively slow moving, do not fly, and may spend days attached to a host, making repellency efforts difficult to quantify. The traditional strict definition of repellency wherein the arthropod makes an oriented movement away from the AI source is therefore not always appropriate in evaluation against ticks, despite being a primary focus and metric in arthropod repellent research [14]. In mosquitoes, repellency is characterized by action that prevents landing on a host. In ticks, repellency can be demonstrated by actions that prevent movement onto a host, direction to a favorable site of feeding, and attachment. Research on chemical repellents against ticks has so far focused on contact repellency. There is a critical need to evaluate and understand spatial repellency of volatile repellent compounds and their potentials for use in personal protection against tick bites.

Herein we present a novel assay to evaluate spatial repellency in ticks, considering the controlled release of an AI, i.e. metofluthrin, transfluthrin, nootkatone, and DEET, against each of the three predominant species of ticks affecting North America: *Dermacentor variabilis, Amblyomma americanum*, and *Ixodes scapularis*. This assay has been named Vertical Tick Assay for Evaluation of Spatial Repellents (VTA-ESR) and provides a quantitative model of ambushing ticks in an evaluation of metrics related to innate host-seeking tick behavior.

## Materials and methods

### Chemicals

Four repellent compounds were tested in this study, including DEET (30% commercial formulation; Ben’s, Littleton, NH, USA), metofluthrin (generic supplier), transfluthrin (Bayer Corporation, Pittsburgh, PA, USA), nootkatone (Sigma-Aldrich, St. Louis, MO, USA). Isopropyl alcohol (IPA, Sigma-Aldrich, St. Louis, MO, USA) was used as a solvent to make test 30% test solutions of metofluthrin (v/v), transfluthrin (v/v), and nootkatone (w/v).

### Ticks

Specific pathogen-free *Amblyomma americanum, Dermacentor variabilis*, and *Ixodes scapularis* adult, female ticks were obtained from the tick-rearing facility at the Oklahoma State University, Department of Entomology and Plant Pathology, National Tick Research and Educational Resource. Additional ticks were flagged from North Amherst, Massachusetts. Ticks were stored in 48 mm (h) x 20 mm (w) plastic vials with plastic caps of 5 mm diameter orifice covered with woven, cotton cloth to prevent ticks from escaping. Four ticks were stored in each vial at 4°C. The containers were removed from refrigeration weekly and opened for 10 minutes. The cloth, vials, and containers were checked thoroughly at this time for evidence of fungal growth. If evidence of growth was noted on the vials, the ticks were moved to clean vials. Ticks were handled with autoclaved forceps and paintbrushes. Two hours preceding the repellency trial, ticks were equilibrated in an incubator at 23°C at 90 % relative humidity (RH). A total of 267 total female ticks were used in this study, including 81 *A. americanum*96 *D. variabilis, and* 90 *I. scapularis*

### Experimental setup

The tick behavior test system consisted of (1) a chemical-emanating device, which was specifically designed for the sustained spatial release of test repellent formulations (Figures 1,2), (2) a tick behavioral test chamber, and (3) a computer-based tick movement tracking system (Ethovision, Noldus Information Technology, Leesburg, VA, USA) (Figure 2). The tick behavioral test chamber was assembled from six clear, acrylic sheets: four 60 cm (l) x 30 cm (w) x 0.5 cm (z) sheets form the bottom, top, front, and back faces, while two 30 cm (l) x 30 cm (w) x 0.5 cm (z) sheets form the sides. The sheets were connected by living hinge plastic connectors (6 cm (l) x 2 cm (w) x 4 cm (z)). Two hinge connectors were used on each face, positioned one inch from either edge (Figure 2). The top sheet rested on top of hinge connectors but were not connected to allow for placement of the active ingredient, and introduction of the ticks into the trials. There was a 5 mm wide opening lining the exterior of the box, created by the placement of the hinge connectors. Between trials, the acrylic sheets were disconnected, cleaned with IPA, and allowed to dry. Three sticks, 32 cm (l) x 0.3 cm (diameter), were used for each climbing experiment. The sticks were adhered to the inside of the top sheet by a 1 cm x 1 cm square of Crayola air-dry clay, cut with a sterile #10 scalpel. The three sticks were placed along the center width of the lid, 2.5 cm from either side and in the center. After each trial, the climbing sticks and clay were discarded, and the walls of the chamber were cleaned with IPA.

**Figure 1.**
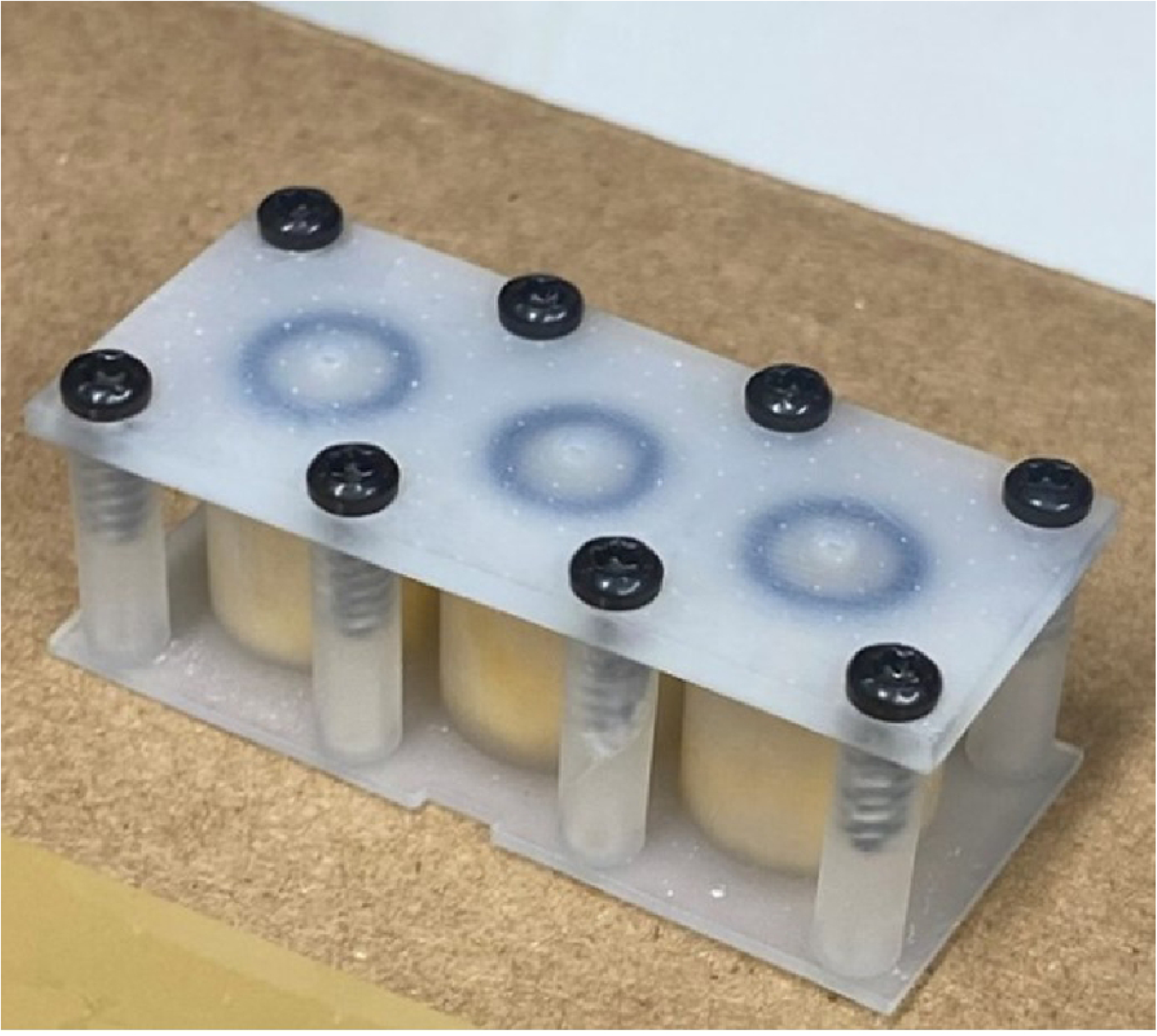
Controlled Release Device (CRD). Diffusion occurs through a small pore shown in the outer surface.

**Figure 2.**
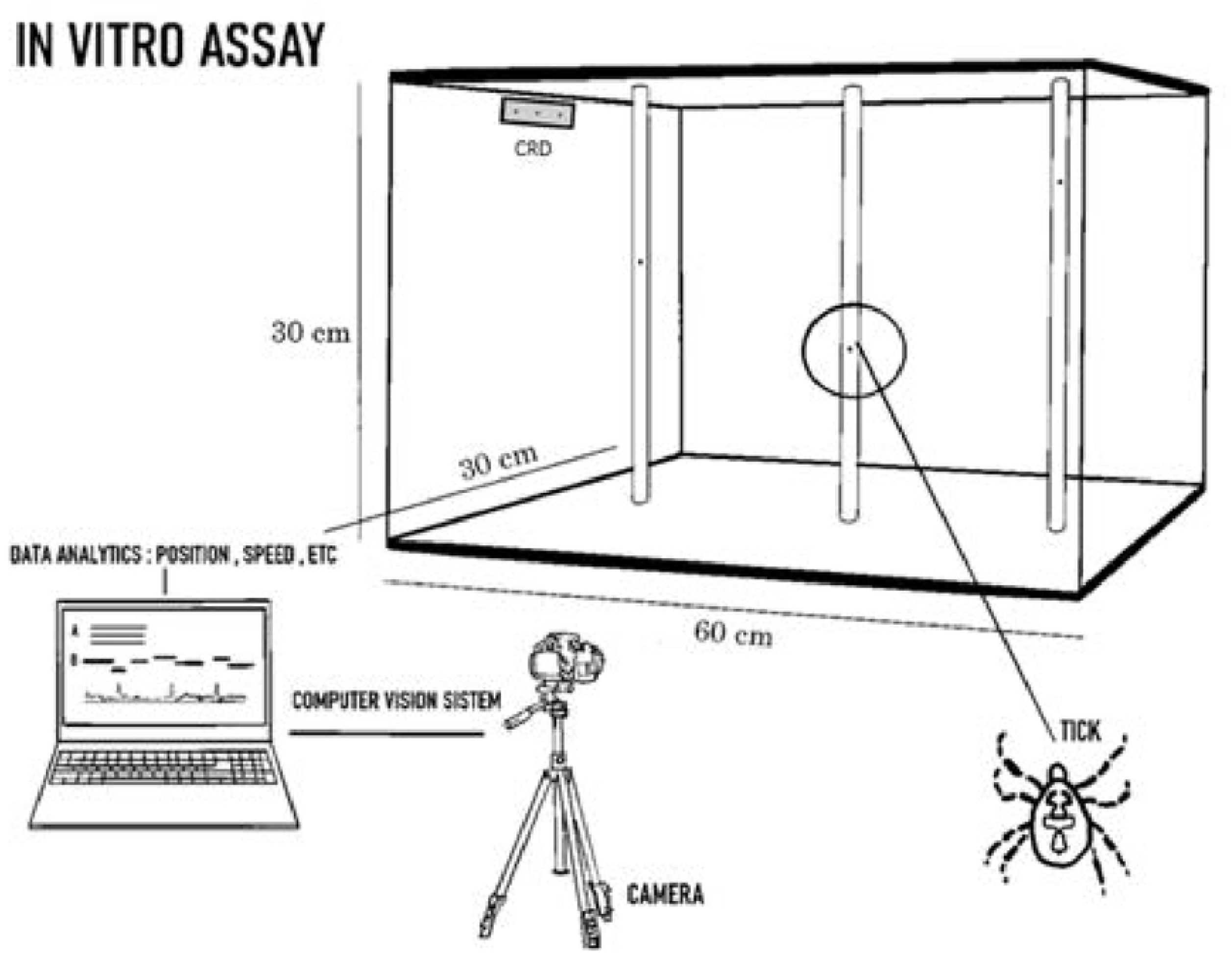
Experimental setup for trials from the perspective of the position-tracking camera. The three 30-cm sticks (fiber diffusers, Simoutal brand) are shown adhered to the top lid with air-dry clay. The device is positioned on upper left side wall in all trials, in the center of the z axis and at the height equal to the top of the stick. The device emits the AI in the direction of the sticks. Clear acrylic sheets (four 30 × 60 cm, two 30 × 30 cm) compose the walls of the box, attached by plastic hinge connectors.

### *In Silico* Simulations

AI release rates from CRDs were characterized using analytical formulations of transfluthrin and metofluthrin, assessing volatilization for a period equal to the duration of the experiment. For this measurement, emanating devices were placed in a sealed bottle and a 50 mL air sample was extracted via syringe chemistry tools based on Gas Chromatography Mass Spectroscopy Analysis (GCMS). Extracted air samples were dissolved in 5 mL of IPA and injected directly into the GCMS for the measurement of AI concentrations relative to a known standard.

To address AI concentration effect on tick behavior, numerical simulations based on Computational Fluid Mechanics (CFD) were performed to address the transport of formulated AI evaporated into still air inside the test chamber. Fluid natural convection, chemical diffusion and gravity momentum were considered. For further detailed description on the simulations, please refer to the Appendix.

### Repellency bioassay

The CRD containing a particular test repellent solution was placed at the upper end of the chamber (Figure 2). An induction time of 20 minutes after each initial device activation was set prior to the introduction of the ticks for each trial. Each trial included 3 female ticks of the same species. One tick was placed at the base of one of the 3 vertical sticks. The array of three climbing sticks in the chamber allowed the assessment of effect of the AI diffusion in the chamber. From the eye of the camera, the CRDs were placed in the upper left hand of the chambers. Hence the three sticks left to right were positioned along a concentration gradient. Ticks were evaluated for inclusion in the trial by briefly placing them at the base of a stick. If the tick climbed the stick, it was included in the trial. Ticks unable to hold on to the stick, detached, or unwilling to climb were excluded. Climbing trials were conducted by removing the chamber lid with the attached climbing sticks, inverting it, placing ticks on the top quarter of the climbing stick and allowing them to climb to the top of that stick. The entire lid with three climbing sticks with ticks at top was then inverted and placed on the walls of chamber so that the ticks were at the bottom of the climbing sticks. These treatments were compared to controls performed in same fashion without AI.

### Video tracking of tick movements

Tick mobility was tracked with a computer vision system [15]. Each experiment was generated with a pre-defined template held consistent through each trial, maintaining constant capture rate, arena centering and size (camera field of view), and detection criteria for tracking. Tick movements were tracked for time periods of 10 minutes. To measure repellency, climbing trials were conducted in presence of AIs and compared with tick activity in absence of the AI. Height (cm) was recorded frame by frame. This allowed the analyses of velocity, displacement, and detachment. Velocity is measured as the rate of movement (cm/sec). Displacement is total distance moved through course of the trial. Detachment is defined as when ticks fall from sticks. An integrative measure of pseudo-questing considered the time ticks spent at the top of the box (27-30 cm) to simulate questing behavior of ambushing ticks. Finally, climbing height reduction considered the cumulative amount of time that ticks spent 27-30 cm, normalized to control trials.

Each trial or experiment, as described above to test responses of each tick species to each compound, was repeated 4 to 6 times to allow statistical analysis of data. Responses to an AI- free environment were used as negative control to allow assessment of effects of the test repellent compounds.

### Statistical analyses

Statistical analyses were performed in SPSS for Windows, version 28.0 following the retrospective correction of insignificant movement captured by the computer vision system [16]. This was done in SPSS through filtration of movement points less than 0.05 cm/s, a point slightly less than the minimum velocity that ticks were observed to move. Measured parameters did not assume the normal distribution. Skewed data were not transformed and were therefore analyzed using non-parametric methods. Mann-Whitney U (MWU) tests were applied to continuous data considering mean velocity and pseudo-questing tendency. All ticks were included in each analysis except for velocity. Ticks that did not move during the trials were omitted from the velocity analysis. Effect size (r) was calculated to provide an indicator of the magnitude of difference between treatment and control groups to supplement probability values. Results were considered significant in cases where U < U_crit_ at a significance threshold α < .05, following SAMPL guidelines [17,18]. These results were then interpreted according to Cohen’s classification of effect size at 1 degree of freedom: 0.1-0.3 small, 0.3-0.5 medium, and > 0.5 large [19]. Two-tailed fisher’s exact tests analyzed the difference in proportions of treatment trials to control trials in climbing success and detachment analyses. Effect size was also calculated for each Fisher’s exact test (φ) and interpreted similarly to those of MWU tests. Significant probability values are reported in figures 6-8 and are presented in tiers: * p < .05, ** p < .01, *** p < .001.

## Results

### Controlled release device characterization and *in silico* model

GCMS results for the first 30 minutes were calculated as 3.1 mg/hr for the transfluthrin formulation and 4.4 mg/hr for the metofluthrin formulation. Concentration plots of transfluthrin and metofluthrin, in parts per billion (ppb), were plotted for 25 minutes in logarithmic scale (Figure 3). A logarithmic concentration plot of transfluthrin and metofluthrin on the sticks is also shown at 25 minutes (Figure 4). Higher AI concentrations were found on the bottom surface.

**Figure 3.**
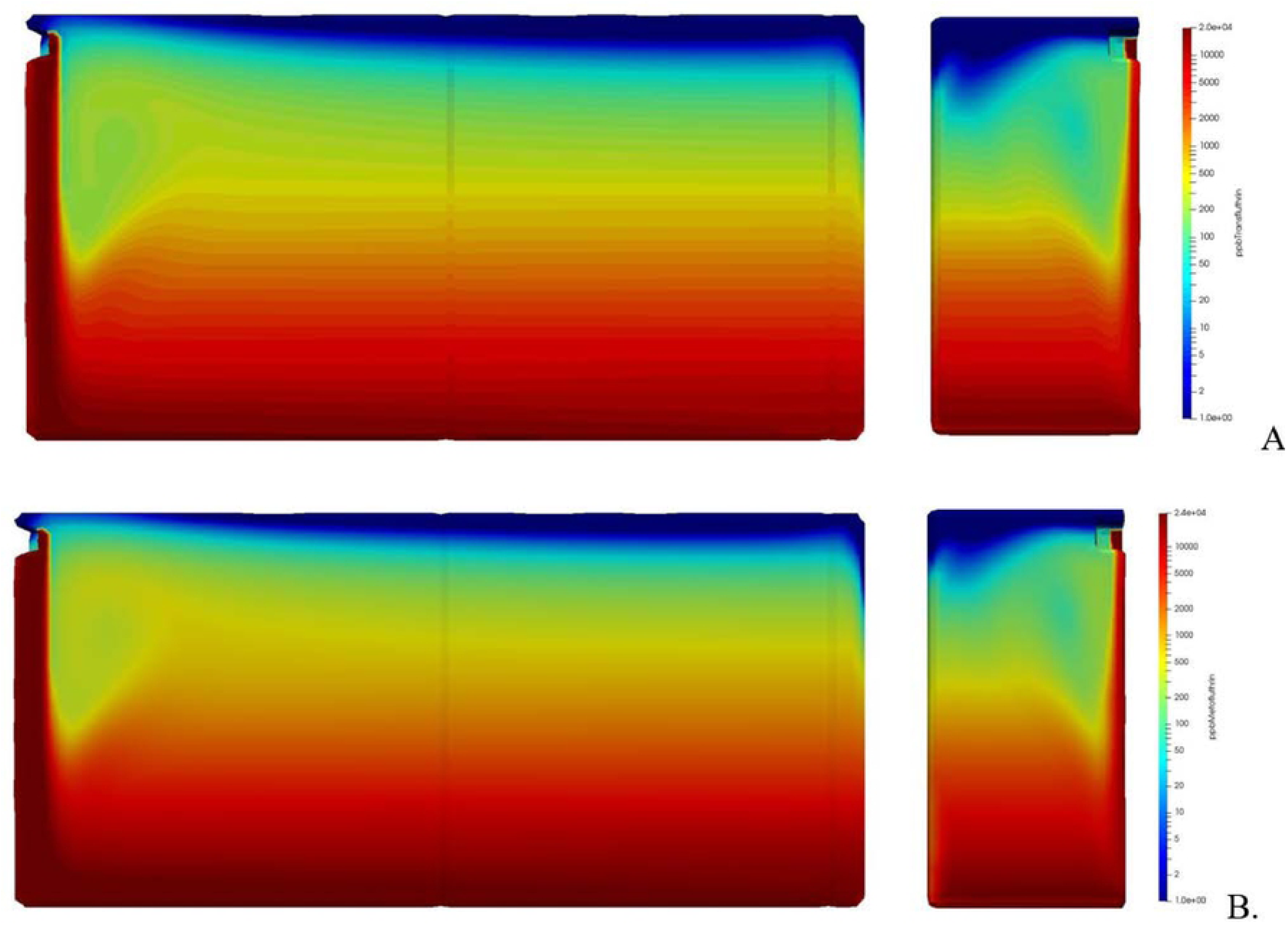
Simulation Results. A. Concentration of transfluthrin in ppb at 25 minutes. B. Concentration of metofluthrin in ppb at 25 min. The larger box represents the full front view of the chamber. The right, smaller portion is a half section from the perspective of the side.

**Figure 4.**
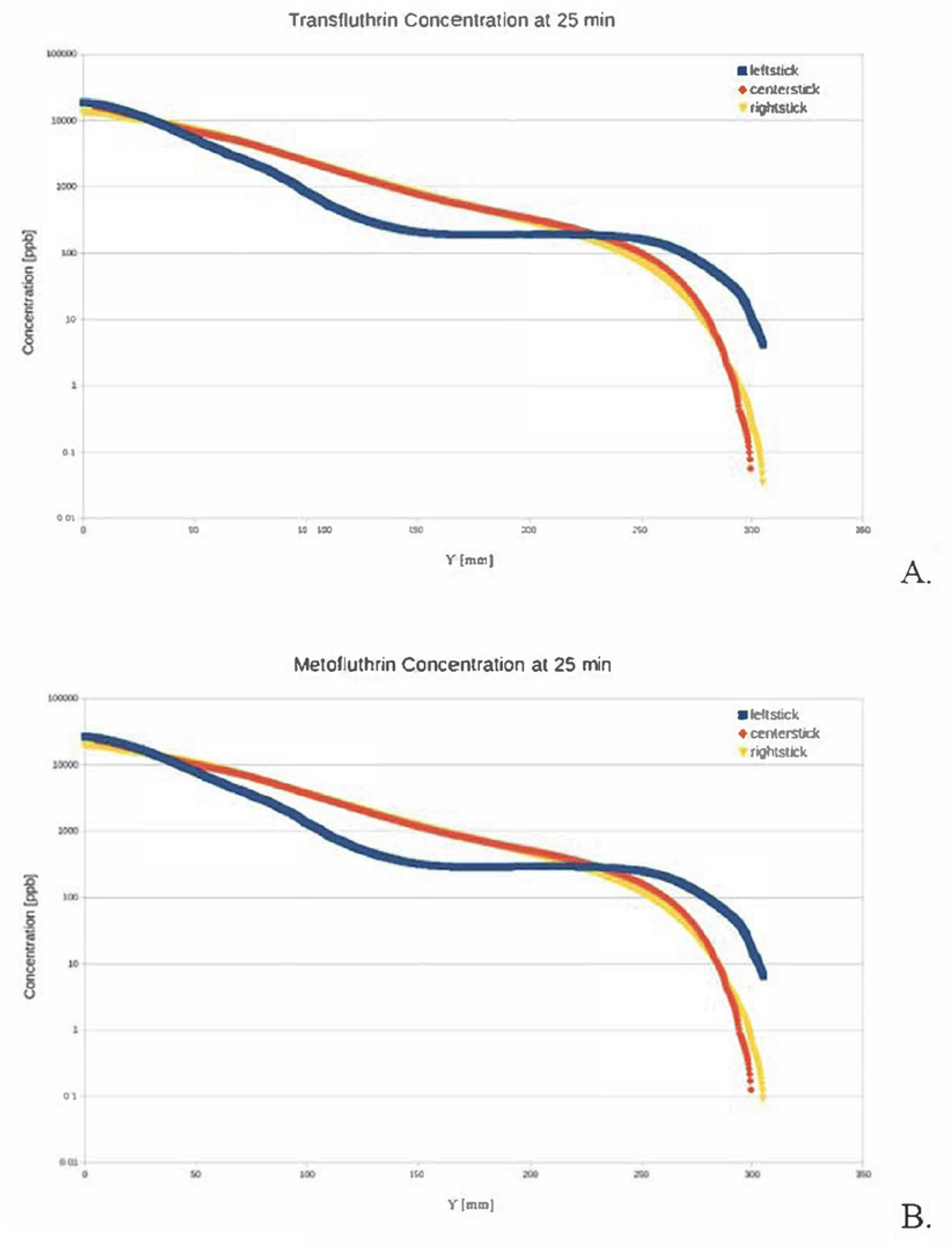
Concentration of transfluthrin (A) and metofluthrin (B) profiles at the sticks at timestamps of 25 min.

This is caused by gravity that dominates the flow, making it plummet since the AIs are heavier than air. Therefore, the fluid density is higher where the AIs and isopropanol are, keeping the concentration levels high on the bottom and low on the top. In the long term however, the concentration increases from the bottom up due to diffusion, as there is a large concentration gradient between the bottom and the top of the container. At higher heights, the concentration remains low, both because gravity tends to keep the heavier molecules at lower heights and diffusion is low because of the low concentration gradient present.

## Control observations

Control trials established baseline behaviors that were compared with ticks exposed to each active ingredient. When the ticks were placed in the assay box, they climbed to the top of their stick. They rarely moved down the stick or detached. *A. Americanum* ticks took slightly longer to orient than the other species at the bottom of their stick but climbed to the top with a mean velocity much greater than that of any *D. variabilis* or *I. scapularis* ticks. They also showed more aggressive behavior, characterized by efforts to attempt to escape the experimental area through the top of the box, breaching the intersection of the stick and the top sheet of the box. Some ticks (2/15) detached, but only after having reached the top. No *D. variabilis* or *I. scapularis* ticks detached. *I. scapularis* ticks move slower, tending to settle slightly below the top of the stick (Table I). The ticks of the same species move at a similar velocity that does not vary much based on time or height. All ticks survived both control and AI trials.

**Table 1.** Comparison of tick detachments among three tick species in response to the control and AI trials.

| Species | Proportion of ticks that detached during trials |  |  |  |  |
| --- | --- | --- | --- | --- | --- |
|  | Control | Metofluthrin | Transfluthrin | DEET | Nootkatone |
| *A. americanum* | 2/15 | 3/15 | 2/15 | 3/18 | 2/18 |
| *D. variabilis* | 0/15 | 4/24 | 2/21 | 1/18 | 2/18 |
| *I. scapularis* | 0/18 | 6/18 | 1/18 | 5/18 | 2/18 |
| Sum | 2/48 (4.2%) | 13/57 (22.8%)<br>P = .010 | 4/54 (7.4%)<br>P = .681 | 9/54 (16.7%)<br>P = .056 | 6/54 (11.1%)<br>P = .276 |

### Tick detachment

Few *A. americanum* detached from their sticks in control trials (2) however only once reaching the top, and no *I. scapularis* or *D. variabilis* in theirs. (Table I). Metofluthrin exposure was associated with the greatest number of ticks detaching in each species in all AI trials (*A. americanum:* (3/15) *D. variabilis:* (4/24), *I. scapularis* (6/18). There was a significant difference between the total proportion of ticks that detached when exposed to metofluthrin (22.8%) compared to controls (4.2%), *P = 0*.*010*. Transfluthrin and nootkatone did not significantly affect the proportion of ticks detaching in their presence (1-2 in each case per species). In the presence of DEET, *D. variabilis* did not show an increase in detachment (1/18). The proportion of *A. americanum* and *I. scapularis* that detached in its presence were both higher (*A. americanum:* 3/18, *I. scapularis:* 5/18), though DEET results were also non-significant.

### Climbing height reduction

The tick response to each AI as measured by position, is plotted showing the mean time ticks spent at positions along the stick during the trials as a measure of height reduction (Figure 5). Metofluthrin resulted in the greatest deterrence in *A. americanum* (74%) and *D. variabilis* (83%) but was slightly less effective in deterring *I. scapularis* (53%). Exposure to transfluthrin showed a similar pattern in each species but was slightly less effective in each case (*A. americanum*: 67%, *D. variabilis*: 82%, *I. scapularis*: 49%). Nootkatone and DEET were ineffective in deterring the presence of *A. americanum* (20%, 0%), but were more effective in *D. variabilis* and *I. scapularis*, showing a larger deterrence (nootkatone: 75% and 65%, DEET: 69% and 84%.

**Figure 5.**
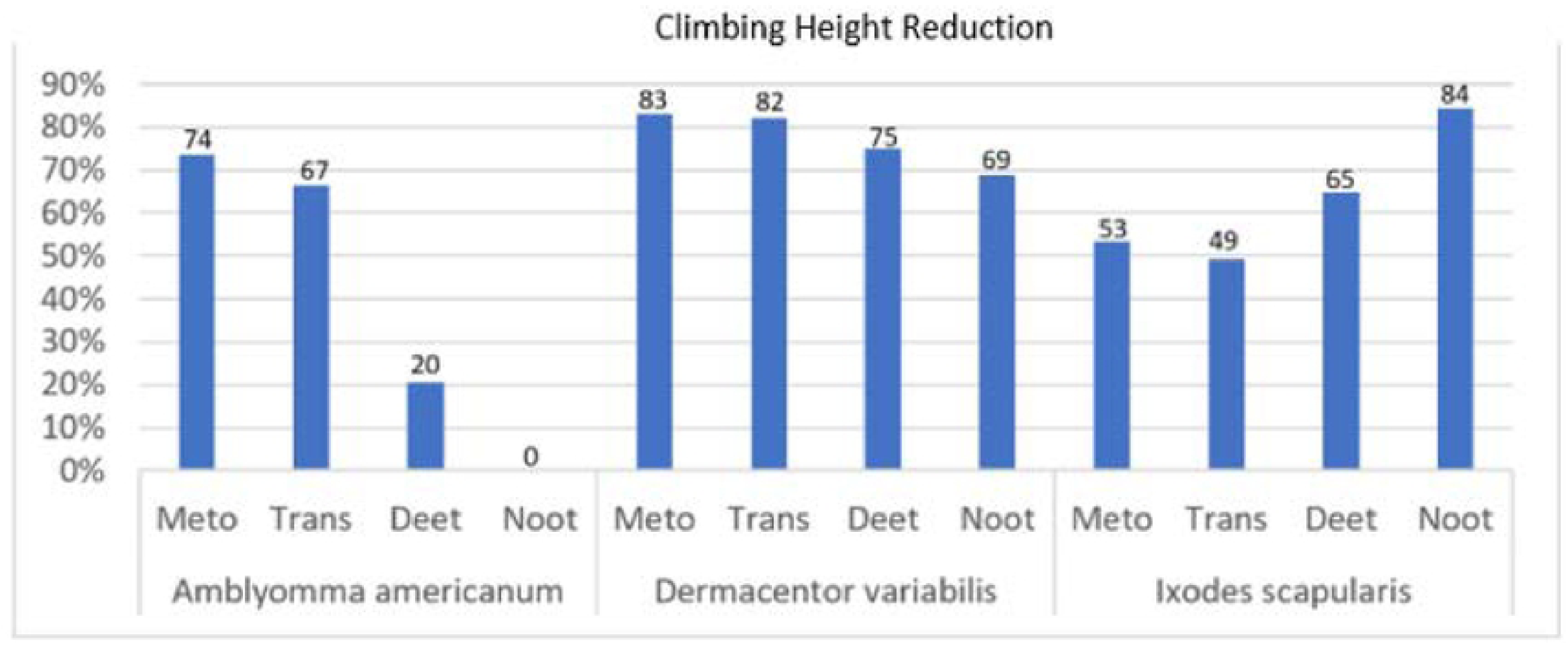
Cumulative distribution of presence 27-30 cm of AI trials are plotted against their respective controls. Transfluthrin and metofluthrin were particularly effective against *D. variabilis* and *A. americanum* but showed less of an effect against *I. scapularis. Nootkatone* and DEET did not affect *A. americanum*. However, they showed larger effects in *D. variabilis* and *I. scapularis*.

### Mean displacement

The mean displacement of control ticks was compared to AI trials (Table 2). Control ticks of each species move approximately one length of the stick by the conclusion of the trial. *D. variabilis* controls moved a mean of 25 cm, *A. americanum* moved 28 cm, and *I. scapularis* moved 29 cm. Metofluthrin resulted in a reduction in displacement for all three species (*D. variabilis:* 11 cm, *A. americanum:* 7 cm, *I. scapularis*: 19 cm). Transfluthrin also resulted in a large reduction of displacement in *A. americanum* (6 cm) and smaller reductions in *D. variabilis* (23 cm) and *I. scapularis* (25 cm). DEET trials resulted in a small increase in displacement in *D. variabilis* trials (27 cm) and a very large increase in *A. americanum* (70 cm), however *I. scapularis* displacement however was reduced (18 cm). Mean *D. variabilis* displacement in the presence of nootkatone was slightly higher than that of the controls (29 cm). *A. americanum* mean displacement was reduced slightly (24.5 cm), while *I. scapularis* mean displacement was increased (52 cm).

**Table 2.** **Comparison of** mean displacements among three tick species in response to the control and repellent compounds.

|  | *D. variabilis* |  | *A. americanum* |  | *I. scapularis* |  |
| --- | --- | --- | --- | --- | --- | --- |
|  | Mean Displacement (cm) | Standard Deviation | Mean Displacement (cm) | Standard Deviation | Mean Displacement (cm) | Standard Deviation |
| Control | 25.30 | 4.41 | 27.75 | 15.18 | 29.02 | 17.54 |
| Metofluthrin | 10.89 | 10.61 | 7.11 | 13.62 | 18.59 | 17.38 |
| Transfluthrin | 22.81 | 10.40 | 5.86 | 9.17 | 24.61 | 25.34 |
| Nootkatone | 28.59 | 5.28 | 24.50 | 17.08 | 52.17 | 60.54 |
| DEET | 27.31 | 8.57 | 69.87 | 47.24 | 17.83 | 16.87 |

### Comparisons of tick activity parameters

AIs showed variable influence on the integrative activity parameters measured (Figures 6-8). Most tick activity parameters measured in responses to DEET and Nootkatone were not statistically significant, and effect sizes of those that were significant tended to be lower than what were observed with transfluthrin and metofluthrin. Metofluthrin and transfluthrin showed more significant and large effects from all perspectives against *D. variabilis* and *A*. americanum but were generally less effective against *I. scapularis* (Table 3).

**Figure 6.**
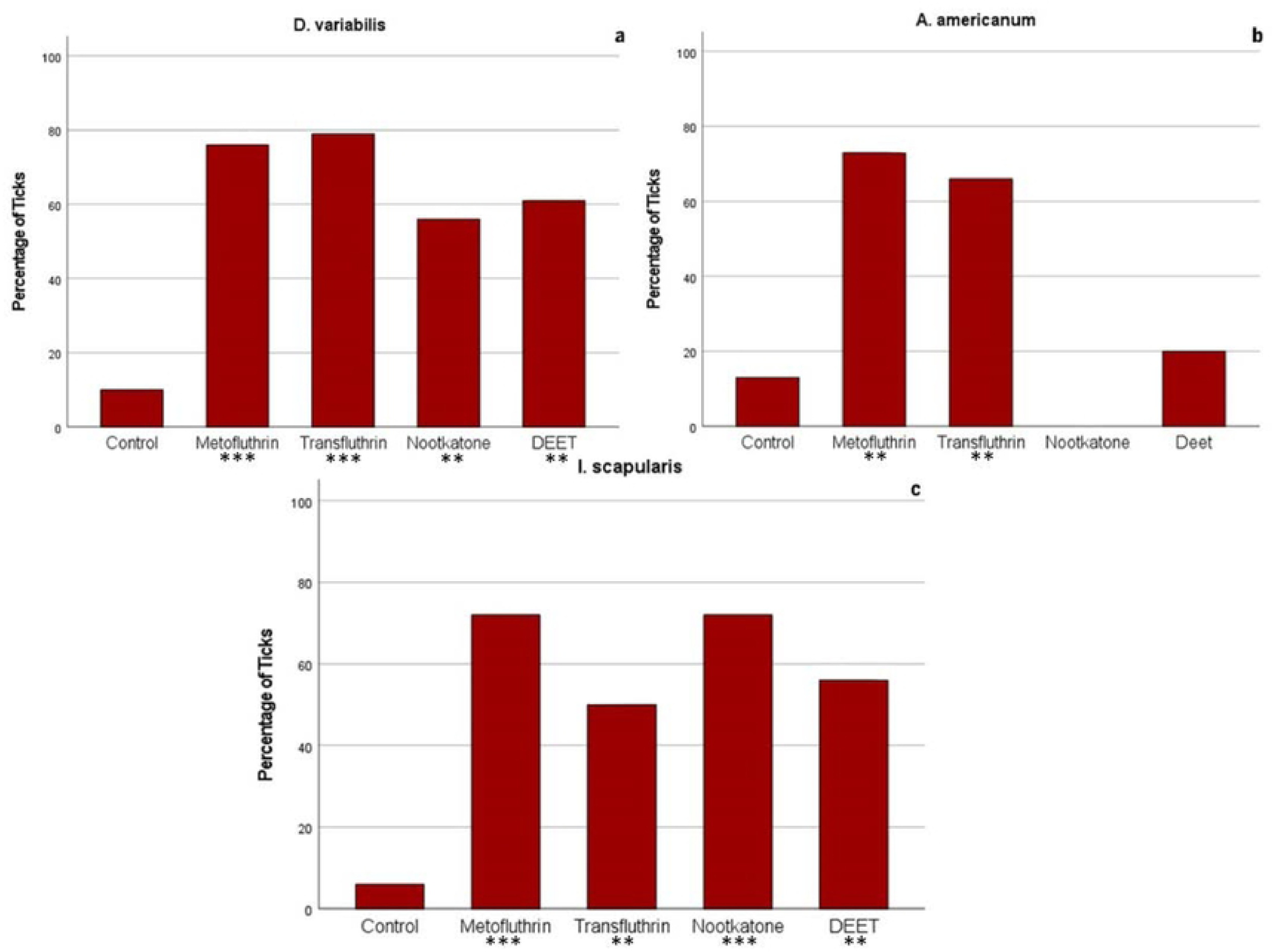
Climbing success of ticks in each control group is compared with ticks in AI trials as a measure of inhibition. A tick that is considered “successful” reaches the 27 cm-30 cm height mark of its stick without detaching and maintains a meaningful presence here, as measured by presence at this point at trial end (t = 600 sec). Difference of proportion show significant differences between AI and controls. *D. variabilis*: Metofluthrin φ = 0.69 (p < .001), transfluthrin φ = .71 (p < .001), nootkatone φ = .52 (p = .004), DEET φ = .56 (p = .003). *A. americanum*: Metofluthrin φ = .61 (p = .003), transfluthrin φ = .58 (p = .002), nootkatone φ = .27 (p = .212), DEET φ = .13 (p = .589). *I scapularis*: Metofluthrin φ = .68 (p < .001), transfluthrin φ = .50 (p < .007), nootkatone φ = .68 (p < .001), DEET φ = .54 (p = .003). Significant probability values are considered in tiers: * p < .05, ** p < .01, *** p < .001.

**Figure 7.**
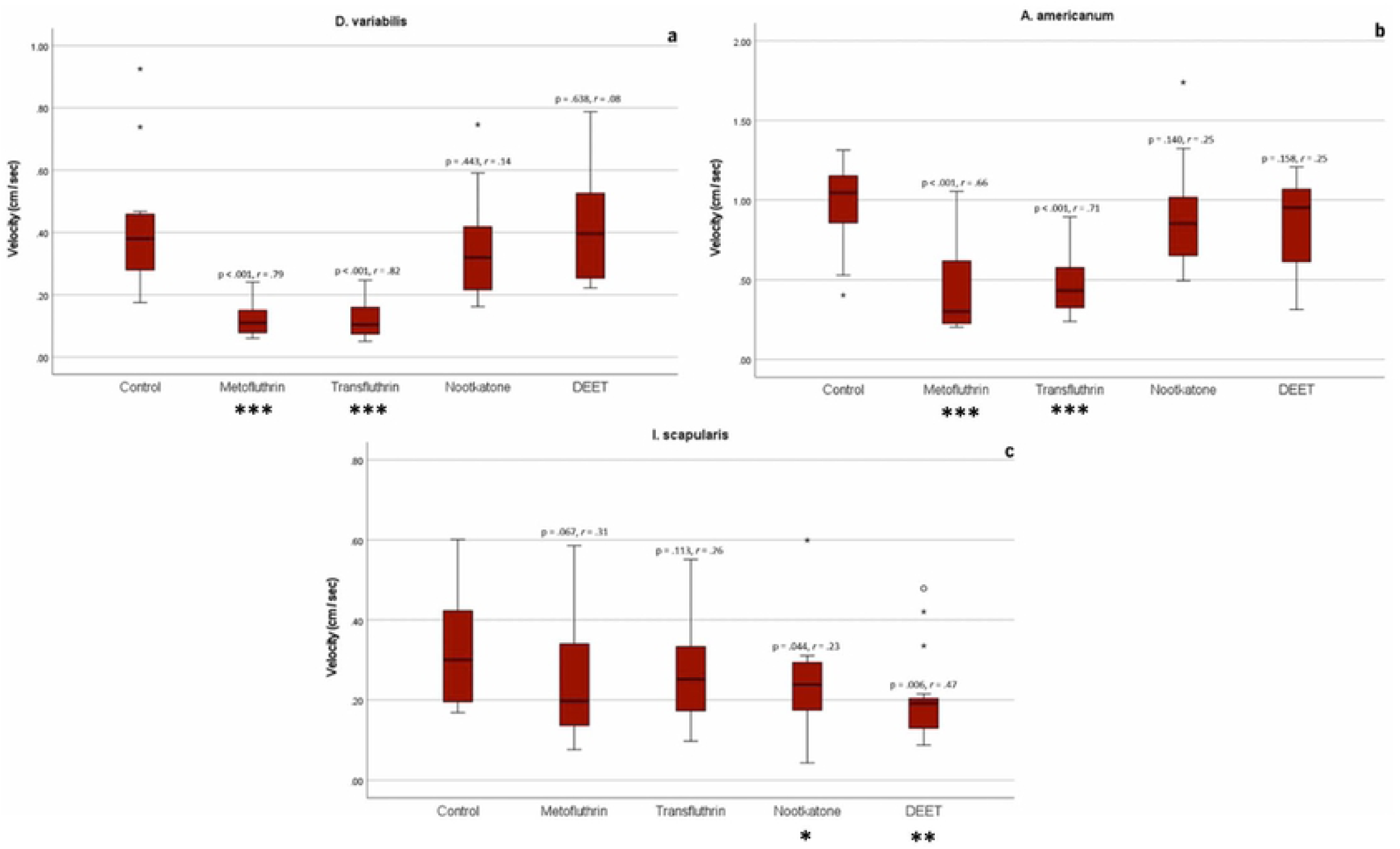
Mean velocity of ticks while moving is compared between AI and control groups, measured in cm/sec. Mann-Whitney U tests showed large, significant differences in *D. variabilis* with metofluthrin (r = .79, p < .001) and transfluthrin (r = .82, p < .001), and much smaller, non-significant differences with nootkatone (r = .14, p = .443) and DEET (r = .08, p = .638). *A. americanum* showed similar results: metofluthrin r = .66 (p < .001), transfluthrin r = .71 (p < .001), nootkatone r = .25 (p = .140), DEET r = .25 (p = .158). *I. scapularis* did not show significant effects with metofluthrin (r = 31, p = .067) or transfluthrin (r = .26, p = 113). Significant differences, though smaller in magnitude, were observed with nootkatone (r = .23, p = .044) and DEET (r = .47, p = .006). Significant probability values are considered in tiers: * p < .05, ** p < .01, *** p < .001. * Outlier of magnitude 1.5-3x IQR ° Outlier of magnitude 3x IQR or greater

**Figure 8.**
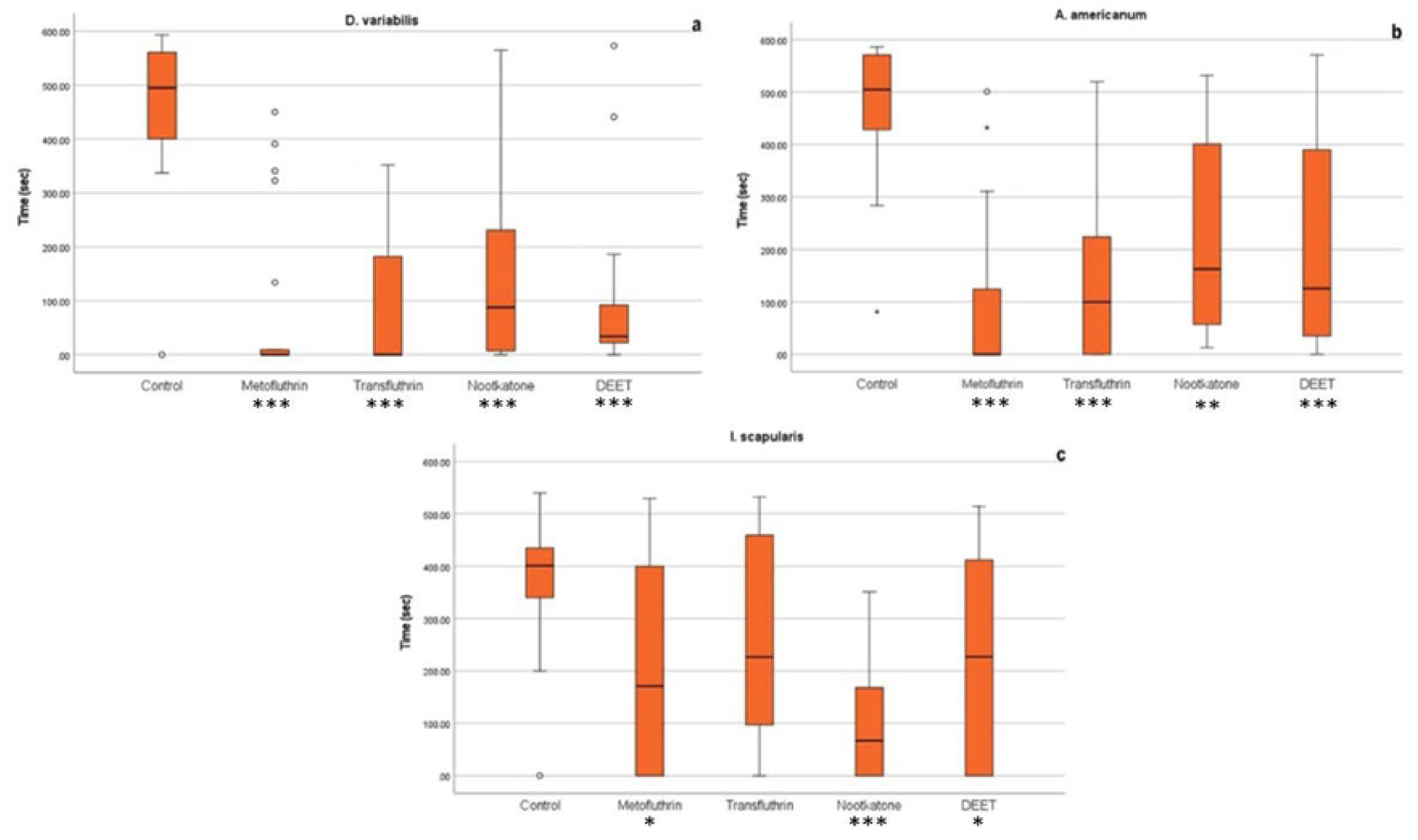
A comparison of pseudo-questing tendency between AI and controls is shown. *D. variabilis* showed significant reductions in the presence of all AIs: metofluthrin r = .76, (p < .001) transfluthrin r = .76, (p < .001), nootkatone r = .63 (p < .001), DEET r = .64 (r < .001). b) *A. americanum* showed similar but slightly weaker results with each AI: metofluthrin r = .76, (p < .001) transfluthrin r = .63, (p < .001), nootkatone r = .56 (p = .001), DEET r = .60 (r < .001). *I scapularis* showed significant results with nootkatone (r = .74, p < .001), DEET (r = .41, .014) and metofluthrin (r = .40, r = .016). Transfluthrin results were not significant (r = .26, p = .121). Significant probability values are considered in tiers: * p < .05, ** p < .01, *** p < .001. * Outlier of magnitude 1.5-3x IQR ° Outlier of magnitude 3x IQR or greater

**Table 3.** Summary of statistical analysis of quantitative behavioral parameters.

| AI | Species | Climbing success ($\phi$) | Velocity (r) | Pseudo-questing (r) |
| --- | --- | --- | --- | --- |
| Metofluthrin | *D. variabilis* | .69*** | .79*** | .76*** |
|  | *A. americanum* | .61** | .66*** | .76*** |
|  | *I. scapularis* | .68*** | NS | .40* |
| Transfluthrin | *D. variabilis* | .71*** | .82*** | .76*** |
|  | *A. americanum* | .59** | .71*** | .63*** |
|  | *I. scapularis* | .58** | NS | NS |
| Nootkatone | *D. variabilis* | .52** | NS | .63*** |
|  | *A. americanum* | NS | NS | .56** |
|  | *I. scapularis* | .68*** | .23* | .74*** |
| DEET | *D. variabilis* | .56** | NS | .64*** |
|  | *A. americanum* | NS | NS | .60*** |
|  | *I. scapularis* | .54** | .47** | .41* |

## Discussion

The global burden of tick-borne disease is addressed through sustainable and integrative approaches that target live tick populations. Increasing incidence in tick-borne disease prompts the development of new options for chemical protection for humans and animals, necessitating both efficacious formulations of AIs and appropriate systems for their delivery. The next generation of innovation in tick protection aims to build on the shortcomings of the current industry standard and identify methods of protection that may apply to a wider range of zoonotic disease-transmitting vectors. Spatial repellency is a novel concept in ticks, however other zoonotic disease-harboring vectors are currently being targeted through volatilized compounds delivered by CRDs and passive methods, greatly contributing to the tactics available in integrative vector management. Metofluthrin and transfluthrin, for example, have demonstrated effective protection from mosquito bites in volatilized formulations [20]. An extension of use into tick control would prove invaluable in providing variety in the ways that ticks can be targeted to reduce the burden of bites and subsequent disease transmission. With applications in regions with overlapping presence of multiple vectors, reduction of disease prevalence from multiple species of arthropod vectors can achieved with single modes of action.

Transfluthrin and metofluthrin were evaluated in the present study alongside two compounds traditionally used in non-volatilized, contact control tactics: the industry standard, DEET, and nootkatone – an acaricidal compound found in grapefruit skin used in environmental sprays for tick control [21]. There is no current standard for assessing spatial repellency in ticks, however the two targets of repellents are defined by the prevention of movement across a “protected” surface and preventing attachment for subsequent feeding and disease transmission. The VTA- ESR assay considers these in analyses of behaviors that are integral to a tick’s successful navigation around these measures, revolving around successful climbing, which is required of a tick for host-seeking and feeding.

Ticks have a finite amount of energy and moisture available to fuel host-seeking. Thus, they must use this supply wisely [22]. In conditions conducive to host-seeking, they climb foliage and passively await a host. Ticks in the control groups for each species reliably climbed to the very top when placed at the bottom of their sticks. They tended to stay at the top, either attempting to escape the box through the top or settle at the top of the stick in a pseudo-questing position. Exposure to all four AIs was associated with significant reductions in pseudo-questing tendency in *D. variabilis* and *A. americanum*. This association was strongest with metofluthrin and transfluthrin in both species. In *I. scapularis*, nootkatone showed the strongest effect, however DEET and metofluthrin showed smaller, significant reductions. Only transfluthrin was not associated with a significant reduction. The deterrence from remaining at this pseudo- questing position may have implications to an inhibition of natural questing in ambushing ticks, however these metrics are unable to make this distinction from other stages of host-seeking and feeding as performed.

In addition to observing gross behaviors as a simulation of host-seeking, an activity analysis of velocity and displacement was performed to visualize any specific effects that AI- exposure may have had on their capability or desire to move, translating to a physical ability to carry out host-seeking and on-host movement. There were several occurrences of large changes in the distance ticks traveled. The greatest of which were with metofluthrin and transfluthrin, which reduced the displacement of all three species. DEET showed a meaningful reduction in *I. scapularis* and an increase in *A. americanum* but did not result in a change in *D. variabilis*.

Nootkatone showed the opposite in *A. americanum* and *I. scapularis*, but also didn’t greatly affect *D. variabilis*.

Metofluthrin and transfluthrin showed very large, significant reductions in mean velocity relative to controls, but were less effective against *I. scapularis*. Nootkatone and DEET were not associated with a change in velocity in *D. variabilis* or *A. americanum* but showed a weak reduction in *I. scapularis* velocity. The reduction in velocity shown by metofluthrin and transfluthrin in *D. variabilis* and *A. americanum* could be evidence of visual effects of AI interference in ticks’ natural ability to move. Increased distance moved relative to controls could indicate deterrence from questing or feeding by keeping the ticks moving. However, a decrease in time and distance moved could also be an indicator of ticks failing to reach a desired location. The changes in tick activity in both velocity and displacement perspectives illustrate effects by the AIs, alluding to applications in repellency evaluation.

Simulation results of transfluthrin and metofluthrin dispersion indicated the formation of a discernible concentration gradient, with greater concentrations distributed towards the bottom of the box and weaker towards the top. Tick natural behavior to climb up was affected by the AI concentration. This effect was visible immediately following tick introduction to the bottom box. Characterized by a propensity away from an immediate climb to the top of the stick (as seen in control trials), ticks in AI groups favored an increased amount of time spent towards the bottom and slower movement where concentrations were highest. The lack of tick movement opposing the concentration gradient indicates that the AIs do not act as a movement barrier at the present concentrations and means of use, but instead immediately disrupt favorable movement patterns aimed at the top of the box, pushing the ticks to continue questing for a safer place. The behavioral change is observed from the beginning of the tick insertion meaning short exposition to AI even in low concentration is enough to disrupt the host seeking will. It is therefore possible that the concentration range used in these trials caused an intoxicating effect that led to a behavioral change.

Detachment is an important indicator of inhibition in the host-seeking and feeding behavior of ticks (Halos 2012). If a tick detaches, it is not feeding or transmitting disease, therefore detaching while moving up the stick could be indicative of a deterrent effect.

Metofluthrin was the only AI that resulted in a larger number of ticks in all three species detach, although DEET exposure was associated in a large proportion of *I. scapularis detaching*.

Transfluthrin and nootkatone were not associated with meaningful increases in detachment relative to controls in any of the three species. Detachment was considered in an integrative metric that also incorporated the height that ticks reached in a success/failure analysis of climbing. Exposure to transfluthrin and metofluthrin was associated with a stronger inhibition of successful climbing, when compared to nootkatone and DEET in *D. variabilis* and *A. americanum*, and showed similar, mild results in *I. scapularis*.

Overall, metofluthrin and transfluthrin showed potential for serving a role in tick protection, generally outperforming nootkatone and the gold standard in today’s commercial tick protection, DEET, in *D. variabilis* and *A. americanum*. Both compounds showed slightly less effect in *I. scapularis*, however comparable to nootkatone and DEET. Nootkatone was particularly ineffective in all metrics when tested against *A. americanum* but performed better in some areas than the other AIs in *I. scapularis*.

Reasoning behind this variation in the degree of differences in behavior observed between species with these AIs is not well-understood. Observed sensitivity of ticks to AIs can vary based on the inherent differences between activity of the ticks, with more active species, like *A. americanum*, generally producing an underestimation of true repellency simply due to their higher speed and agility. Beyond this, however, physiological and molecular differences between species likely result in differences in response.

The basis of tick olfaction begins on the terminal segment of the front legs, within the Haller’s organ [23]. The Haller’s organ is comprised of an anterior pit that detects humidity and a capsule that houses physiologically diverse olfactosensilla. The porous walls of olfactosensilla allow vaporized odorant molecules to enter and reach the lymph. Here, odorant binding proteins are selectively bound by odorant molecules. They are then solubilized and shuttled to odorant receptors on the dendrite of olfactory receptor neurons. Olfactory receptor neuron-reception of host-derived chemical stimuli, such as carbon dioxide, guides the host-seeking and questing process. Dendritic branching increases sensory cell surface area for detection of low concentrations of these odorant molecules, thus allowing this host-detection to occur at a distance [24]. Downstream molecular physiology beyond this has yet to be characterized, however variations in the structure of odorant binding proteins, odorant-degrading enzymes, degree of dendritic branching, and odorant receptor physiology may contribute to the interspecies differences in sensitivity to the active ingredients. Odorant receptor variability is likely similar to observed differences between species of other arthropods. For example, amino acid sequencing has revealed significant variability in mosquito odorant receptor composition [25]. However, genomic investigations into molecular basis of tick olfaction have failed to identify odorant receptors in the genome [26]. Further research is therefore needed to characterize the molecular mode of action in tick olfaction to compliment the analysis of novel tick-targeting chemicals. Because tick behavior is so olfactory-driven, further insight would be useful to guide product development [27].

The current study identified several behaviors that can help investigate effects of volatilized compounds against ticks. *In vitro* methods in preliminary assessments of novel AIs are limited in generalizability to more natural conditions, as the evaluation of repellency for practical use requires an assessment of both the novel AI and its intended formulation in a setting that considers the external factors that may negatively impact product efficacy. Abrasion, temperature, humidity, and wind can affect the potential of a formulation and alter the extent to which the targeted vector responds. Furthermore, factors such as different binding properties to clothing, hair, and skin and trans-epithelial transport can affect the environmental diffusion of the AI [10]. The present study is however an integral early step in the product development process. Ticks have a natural tendency to climb. In the absence of host cues, a demonstration of suppressed efforts to reach desirable positioned, modeled with the vertical climb assay is a first step to determining possible effects. Subsequent studies can build on this work to incorporate more environmental conditions, host cues, and evaluate the AIs at different concentrations, release rates, and in different delivery methods.

## Conclusion

The development of an ideal repellent requires an active ingredient with a formulation that can offer efficacious protection against diverse disease-transmitting vectors in a safe, pleasant formula for consumer use [28]. Applied to ticks, a chemical should operate in two levels of tick protection: preventing travel over a treated surface and preventing attachment. The current assay is unable to distinguish which of the two are being simulated, however the behavior changes that are considered here may be applicable to each. Tick response to volatilized compounds, as opposed to tactile chemoreception, has been speculated in the past but has yet to be effectively demonstrated. The VTA-ESR is therefore useful for the evaluation of several behavior factors applicable to natural tick activity. Exposure to all four AIs was associated with significant changes in tick behavior of varying degree. Transfluthrin and metofluthrin exposures showed an overall greater extent of behavioral differences in all three species. The magnitude of effect for all AIs was reduced in *I. scapularis* when compared to *A. americanum* and *D. variabilis*. This study serves as an initial analysis of spatial repellency in ticks and a preliminary assessment of these AIs for future field application, identifying changes in behavior associated with non-tactile control methods which vary by species. Future studies are needed in the presence of more natural conditions to characterize effects in nature, however the results presented here are integral to reaching this step.

## Acknowledgements

The authors would like to thank Mr. Kevin Smith as representative of Bayer Environmental Sciences for providing the Active Ingredient to conduct the research. His enthusiasm and corporate contribution are greatly appreciated.

This article reports the results of research only. Mention of a proprietary product does not constitute an endorsement or a recommendation by the authors, USDA, or DoD for its use. The USDA is an equal opportunity provider and employer.

## Author contributions

Conceptualization: Elman, Li, Rich, ….. Funding acquisition: Elman, Rich

Formal Analysis: D’hers, Olivera, Roig, Siegel Investigation: Siegel

Methodology: Elman, D’hers, Perry, Rich, Siegel Supervision: Rich

Writing – Elman, D’hers, Rich, Siegel

Writing – review & editing: Elman, D’hers, Li, Rich, Siegel

## Notes

### Competing Interest Statement

The authors have declared no competing interest.

